# Spike protein multiorgan tropism suppressed by antibodies targeting SARS-CoV-2

**DOI:** 10.1101/2021.07.30.454520

**Authors:** Molly Brady, Abigail Combs, Chethana Venkatraman, Alexander Solorzano, Angelique Johnson, Conor McQuaid, Akib Rahman, Hannah Leyva, Wing-Chi Edmund Kwok, Ronald W Wood, Rashid Deane

## Abstract

While there is clinical evidence of severe acute respiratory syndrome coronavirus 2 multiorgan tropism in severely infected coronavirus 19 patients, it’s unclear if there is differential multiorgan biodistribution and organ uptake in healthy young individuals, a group that usually has asymptomatic to moderate coronavirus 19 symptoms. In addition, for antibody therapies and vaccines that target the spike protein, it’s unclear if these reduce severe acute respiratory syndrome coronavirus 2 or spike protein multiorgan tropism equally. We used fluorescently labeled spike protein near infrared fluorescence to study viral behavior, using an *in vivo* dynamic imaging system, in young mice. We found a spike protein body-wide biodistribution followed by a slow regional elimination, except for the liver, which showed an accumulation. Spike protein uptake was highest for the lungs, and this was followed by kidney, heart and liver, but, unlike the choroid plexus, it was not detected in the brain parenchyma or cerebrospinal fluid. Thus, the brain vascular barriers were effective in restricting the entry of spike protein into brain parenchyma in young healthy mice. While both anti-angiotensin converting enzyme 2 and anti-spike protein antibodies suppressed spike protein biodistribution and organ uptake, anti-spike protein antibody was more effective. By extension, our data support the efficacy of these antibodies on severe acute respiratory syndrome coronavirus 2 biodistribution kinetics and multiorgan tropism that could determine coronavirus 19 organ-specific outcomes.

## Introduction

The recent pandemic is caused by a coronavirus (CoV) called SARS-CoV-2, which elicits severe acute respiratory syndrome (SARS), an infectious disease, COVID-19[1]. The virus’ main route of entry into the body is by inhalation of droplets; thus, the disease is manifested as respiratory dysfunction leading to pneumonia in severe cases [2–10]. The virus enters host cells by interacting with facilitators, mainly angiotensin converting enzyme 2 (ACE2) receptor, which is widely distributed in many tissues [10–19] . The spike protein (SP) on the viral plasma membrane (a crown-like appearance) avidly binds with membrane bound ACE2 [20]. However, many non-respiratory organs are also affected, such as the heart [21–23], kidneys [24,25], liver [26–29], and brain [3,30–35], which can lead to multiorgan failure in severe cases of susceptible individuals. However, most of the COVID-19 cases are asymptomatic, mild or moderate [5,36,37]. It’s unclear if SARS-CoV-2 is distributed equally to all organs in these cases or if it’s a consequence of cardio- respiratory failure and/or systemic inflammation as seen in the severe cases. In addition, for antibody therapies, such as anti-SP and anti-ACE2 antibodies, and vaccines that target the SP, it’s unclear if these reduce SARS-CoV-2 multiorgan biodistribution equally. In vaccines, such as mRNA vaccines, the translated SP is released into interstitial fluid/blood, distributed to many organs and triggers an immune response. Thus, we studied the biodistribution of intravenously injected SP and tested the effect of anti-ACE2 and anti-SP antibody on SP regional biodistribution and organ uptake using an external *in vivo* dynamic imaging system and ex *in vivo* tissue analysis.

Herein, we show that SP had a body-wide biodistribution, slow regional elimination, except for the liver, which showed an accumulation, and differential organ uptake. SP uptake was highest for the lungs, and this was followed by kidney, heart and liver, but not in brain parenchyma. SP was present in the choroid plexus, but there were no detectable SP levels in CSF. Thus, the brain vascular barriers were effective in restricting the entry of a viral protein (SP) into brain parenchyma and CSF in young healthy mice. Also, SP was present in the salivary glands. While both anti- ACE2 and anti-SP antibodies suppressed SP regional biodistribution and organ uptake, anti-SP antibody was more effective. The data suggested that these antibodies, especially anti-SP antibody, can effectively reduce the SP biodistribution and multiorgan tropism, and thus, may contribute to the efficacy of these therapies or vaccines.

## Results

### Spike protein is widely distribution and slowly eliminated

First, the biodistribution pattern of a SARS-CoV-2 spike protein (SP-NIRF), after intravenous injections, in 2-3 months old male adult mice was established. The SP-NIRF signal was acquired for the whole mouse in its supine position to simultaneously observe regions that are likely affected by the virus using an external imaging system (**Supplemental Fig.1A**). Thus, for this analysis the region of interests (ROIs) were the neck, thorax, upper abdomen, lower abdomen and paw based on preliminary imaging (**Fig.1A**). We confirmed that NIRF signal can be detected under the rib cage (**Supplemental Fig.1B**), as we reported for the skull bone [39]. Following the injection, SP- NIRF signal was increased to a peak then gradually decreased in all these ROIs, except the upper abdomen, which slightly increased after a trough at about 30 minutes (**Fig.1B**). While there was signal on the whole mouse, including the skin, it was most pronounced for the neck (mainly salivary glands) and the upper abdomen (mainly the liver). To correct for possible experimental variations the data were standardized to the peak signal, and the same patterns for the signal profile were observed at each ROI (**Fig.1C**).

**Fig. 1.**
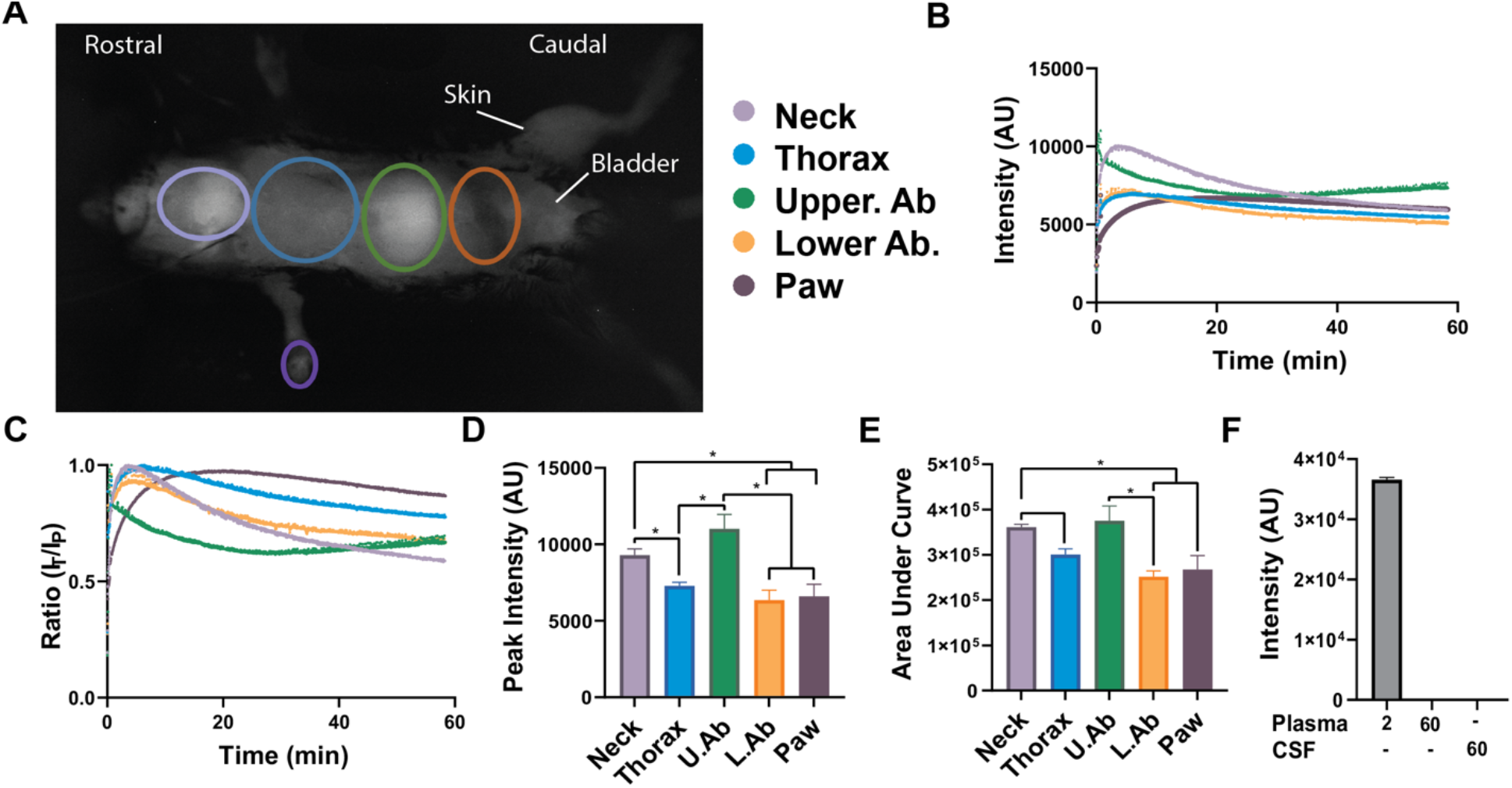
Body-wide biodistribution and slow elimination of a SP in mice. **A**) External in vivo dynamic SP-NIRF image of a mouse after its intravenous injection (after a few minutes). Regions of interest (ROIs) selected for the analysis. **B**) Representative SP-NIRF intensity-time profile for each ROI over 60 min. **C**) Standardization of data by dividing intensities at each time point by the peak intensity (IT/IP ratio). **D**) Peak intensity for the ROIs. **E**) Area under the curve (AUC) for the ROIs. Values are mean ± SEM, N=5 young male mice (2-3 months old). **F**) Plasma intensities at 2 and at 60 min, and CSF levels at 60 min. Values are mean ± SEM, N=3 young male mice. AU (arbitrary units).

The peak SP-NIRF signals were highest for the neck and upper abdomen, which was 1.3- to 1.9-fold greater than that for the thorax, lower abdomen and paw, but there was no significant difference between these regions (**Fig.1D**). While the area under the curve (AUC), the overall regional SP-NIRF exposure, was also similar for the upper abdomen and neck, these were greater by 1.30 to 1.8-fold compared to the thorax, lower abdomen and paw. However, there was no significant differences between the AUC for the thorax, lower abdomen and paw (**Fig.1E**).

The elimination rate constants were no significant for the neck, thorax, and lower abdomen, which had elimination phases (**Supplementary Fig. 1C**). For the upper abdomen, after its distribution the rate of SP-NIRF accumulation was determined from the liner regression of the rising phase of the intensity-time profile (**Supplementary Fig.2D**). SP-NIRF signal was detected in the bladder but this was variable, perhaps, due to changes in urination as this was not controlled (**Supplementary Figure 1E**). SP-NIRF plasma and CSF levels were not detectable at 60 minutes (**Fig.1F**), while the ROI signals were higher suggesting that there was retention within these regions.

In summary, these data show that SP-NIRF was widely distributed within the body of young mice, and taken up in many organs, especially in the upper abdominal (mainly liver) and neck (mainly salivary glands) regions. There was an elimination phase for the neck, thorax and lower abdominal regions, while for the upper abdominal region there was accumulation of SP-NIRF after its distribution, instead of elimination. Peak levels were highest for the neck and upper abdominal regions, which may indicate greater uptake even though there was an elimination phase for the neck region but an accumulation phase for the upper abdominal region. SP-NIRF signals in the paw increased exponentially to a plateau and was almost constant thereafter. By extension, these data suggest that there will be differential multiorgan tropism of the virus in young mice, which could lead to organ dysfunction/failure in susceptible individuals, assuming SP mimics the viral distribution since this is due mainly to SP-host cells interaction.

### SP uptake suppressed by anti-ACE2 and anti-SP antibodies

ACE2 receptor is widely distributed in the vasculature and in tissues, including lungs, kidneys, heart, and possibly the cerebrovasculature [10–12,14,19,32,41,42]. SP interaction with ACE2 receptor would indicate potential viral uptake regions, and anti-ACE2 antibody will decrease viral-host cell interaction. Anti-SP antibody, as used in passive immunity, generated in active immunity, and produced by vaccines, will interact with the SP on SARS-CoV-2 to eliminate the virus, and also reduce viral-host cell interaction. For some of the vaccines, the SP produced by the vaccine will be released from cells and distributed in the body before it can mount an immune response. Anti-SP antibody in the circulation can then interact with the SP on the virus. Thus, we tested the effect of anti-SP antibody and anti-ACE2 on the regional biodistribution, elimination kinetics and organ uptake of a SP to establish if there are differential effects. We confirm that the anti-ACE2 antibody interacts with mouse ACE2 (**Supplementary Figure 2A**).

In separate groups of mice, anti-ACE2 antibody (T1, 10 μg) or anti-SP antibody (T2, 10 μg) was injected 15 minutes prior to the SP-NRF injections. While the general pattern of the ROIs intensity profile curves was similar, the values were lower than that for the young control mice (**Fig. 2A-B**; **Supplementary Fig. 2B-C**; **Fig. 1B**). The peak SP-NIRF signals were significantly higher in the control mice compared to the T1 treated mice, by 1.8-fold for the neck and thoracic regions but not significantly for the upper abdomen, lower abdomen and the paw (**Fig.2C**). In contrast, for T2 treatment it was higher for the controls by 2.0-to 3-fold for all ROIs (**Fig.2C**). The AUC was significantly greater in the control mice compared to the T1 or T2 treated mice by 1.8- to 2.5-fold in all the ROIs, except for T1 and the paw (**Fig.2D**). The intensities were transformed to the Ln scale and the elimination rate constants determined from the disappearance phase of the graph (**Fig. 2E**). T1 and T2 increased the elimination rate constants for the neck by 2.0- to 2.5- fold but not for the thorax and upper lower abdomen compared to controls (**Fig. 2F**). For the upper abdomen, the rate of SP-NIRF accumulation after its distribution was reduced by 5.5-fold with T2 but not by T1 compared to controls (**Fig.2G**). Also, T1 and T2 reduced SP-NIRF distribution (AUC) for the paw (**Fig. 2H**).

**Fig. 2.**
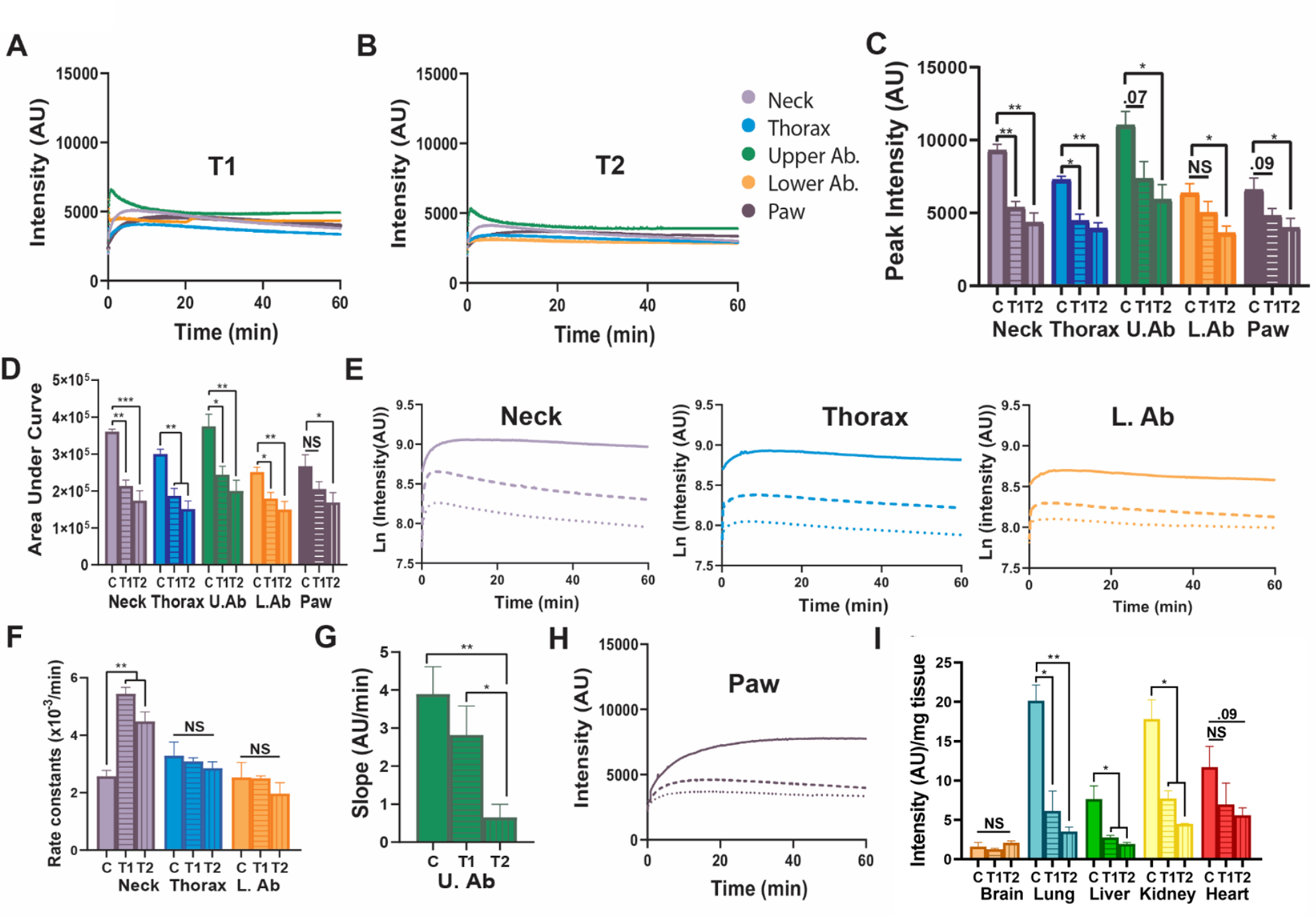
Anti-ACE2 and anti-SP antibodies reduced SP biodistribution. **A-B**) Representative SP-NIRF intensity-time profile at the ROIs for T1 (anti-ACE2 antibody) and T2 (anti-SP antibody) in 2-3 months old mice after its intravenous injection. **C**) Peak intensity for the ROIs. **D**) Area under the curve (AUC) for the ROIs. **E**) Semi-Ln plots for the intensity-time profile of the neck, thorax and lower abdomen ROIs (continuous line (control), dashed line (anti- ACE2 antibody) and dotted line (anti-SP antibody). **F**) Elimination rate constants for the neck, thorax and lower abdomen. **G**) Slopes for the accumulation phase for the upper abdomen. **H**) Intensity-time profile of the paw. **I**) Organ SP-NIRF intensities/mg wet weight at 60 min. Values are mean ± SEM, N=4-5 young male mice. AU (arbitrary units).

Collectively, these data show that both T1 and T2 reduced the biodistribution to the ROIs, with T2 (anti-SP antibody) more effective, in general. Interestingly, for the neck region (mainly the salivary glands, but also include the pharynx and larynx) T2 was effective in increasing the removal of SP-NIRF (higher elimination rate constant). Similarly, T2 was effective in reducing the accumulation phase of the upper abdomen (mainly the liver). For the paw T2 reduced SP-NIRF peak intensity and the AUC. In contrast, for the thorax and lower abdomen, there are many potential sources of the SP-NIRF signals, including lungs and heart for the thorax, and intestines and kidneys for the lower abdomen. Thus, by extension, therapies targeting ACE2 and the SP, and especially vaccines targeting SP are effective in suppressing the biodistribution of the virus to organs in this model.

### Ex vivo organ SP distributions

To better understand which organ was associated with the SP-NIRF, at the end of the experiment, organs (brain, lungs, liver, kidneys, spleen, intestine and spine) were removed, washed in cold PBS and imaged using the same parameters as in the in vivo imaging. There were no significant differences in the intensities between the dorsal and ventral surfaces of the brain even though there are more visible surface blood vessels on the ventral surface (**Supplementary Fig.2D**). Since these organs vary in weight, signal intensities were standardized per unit wet weight. SP-NIRF uptake in the lungs, kidney and heart was 8.0- to 12-fold greater than that of brain (**Supplementary Fig.2E**). The brain had the lowest SP-NIRF uptake while the lungs had the highest. SP-NIRF signals for the spleen was low, but could contributed to the upper abdominal ROI (**Supplementary Fig.2 F-G**). Likewise, the intestine contributed to the SP-NIRF signal for the lower abdomen. Interestingly, SP-NIRF signal for the small intestine was mainly detected in the duodenum (**Supplementary Fig.2 H**). After 60 minutes, there was very low SP-NIRF levels for the salivary glands (**Supplementary Fig.2I**). Other tissues may contribute to signal in the neck region, such as the trachea, tongue and larynx and pharynx. T1 and T2 reduced the SP-NIRF uptake by the liver, lungs and kidney by 4- to 7-fold, but not for heart and brain compared to control young mice (**Fig.2I**). T1 and T2 had no significant effect on the SP-NIRF uptake by the brain. These data show that anti-SP antibodies can effectively reduce SP multiorgan biodistribution, and thus, by extension should reduce the virus effects on these regions.

### Ex vivo analysis of tissue sections

To establish whether the SP was taken up within the organs we analyzed SP-555 uptake and compared it to a reference protein of similar molecular weight (OA-488). Sagittal sections of the head show that most of the SP-555 and OA-488 were present in peripheral tissues, with limited entry, if any, into the brain parenchyma (**Fig.3A**) compared to that of brain sections from non-injected mice (**Supplementary Fig.3A-B**). However, there were SP and OA signals in the region associated with pituitary gland, which lacks a blood brain barrier (BBB), and the choroid plexus, but not across the cribriform plate (**Fig. 3A-C**). Coronal brain sections confirmed the absence of detectible SP-555 signal in brain parenchyma, although there were signals for both SP-555 and OA-488 associated with the choroid plexus compared to samples from non-injected mice (**Fig.3D**; **Supplementary Fig.3B; Supplemental 4A**). There was no significant association of SP with the olfactory bulb, blood vessels or neurons (**Supplementary Fig.4B-D**). In contrast, the spinal cord had detectible signals for both SP-555 and OV-488, compared to samples from non-injected mice (**Fig.3F; Supplementary Fig.3C; Supplementary Fig.4E-F**), possibly due to a greater permeability of the spinal capillaries than that of brain [43]. Thus, the low brain SP-NIRF levels could be due mainly to the choroid plexus and possibly areas lacking BBB, such as the pituitary gland. Thus, the CNS vascular barriers seem to be effective in limiting SP entry into brain parenchyma.

**Fig. 3.**
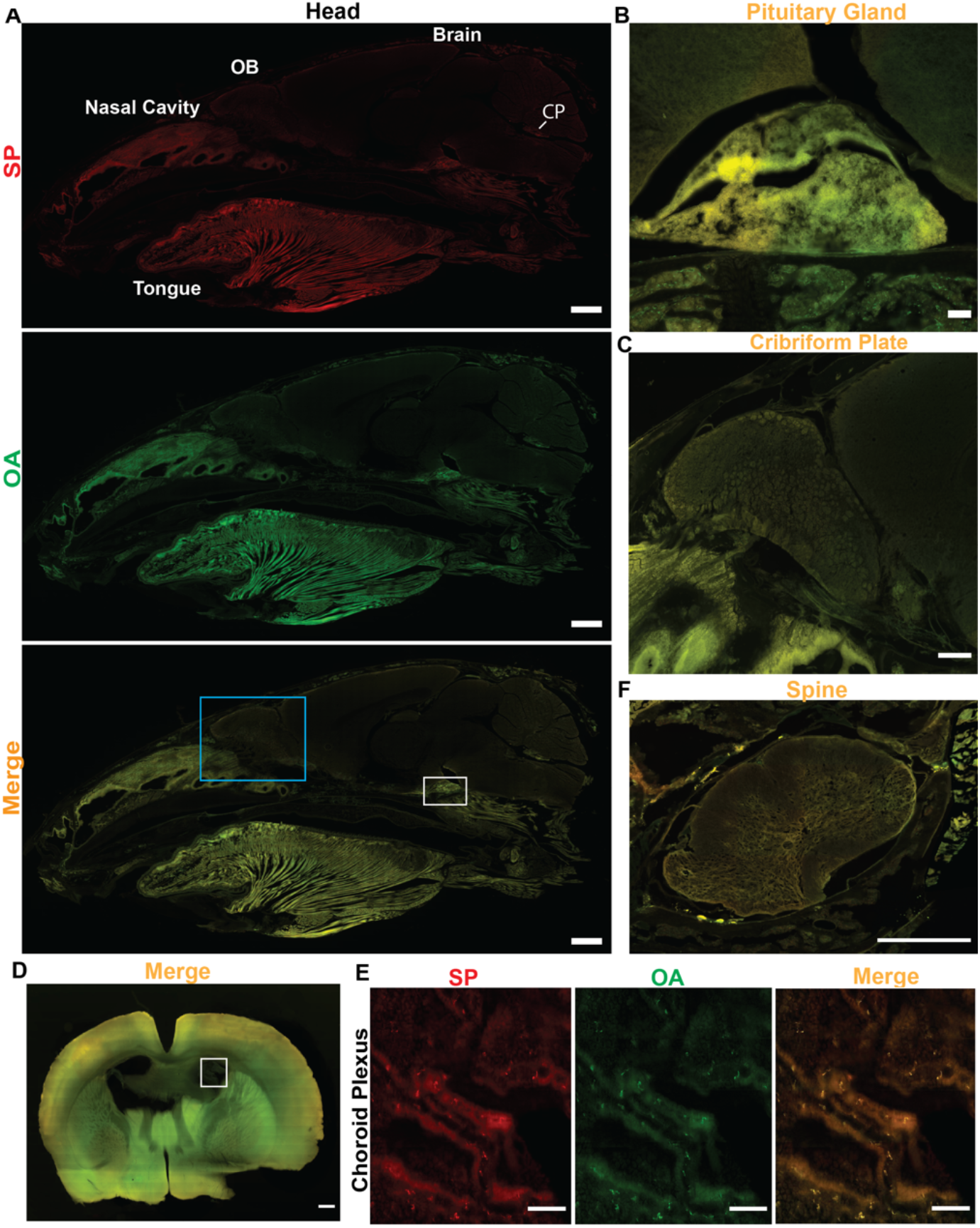
SP is not present in brain parenchyma. **A**) Representative sagittal section of the head for a young mouse showing distribution of SP-555, OA-488 and merged images. OB (olfactory bulb). CP (choroid plexus). **B)** Magnified region (pituitary gland) of the white box in A. **C**) Magnified region (nasal/cribriform plate) of the blue box in A. **D)** Representative coronal sections of the merged images of SP-555 and OA-488. **E**) Magnified region (choroid plexus) of the box in panel D. **F**) Merged image of the spinal cord showing the present of both tracers. Scale bars: A (1mm); B-D (100 μm); E (0.5 mm). Values are mean ± SEM, N=3.

For the peripheral tissues, there were significant distribution of SP-555 and OA-488 in whole section analysis for the lungs, liver, kidney and heart (**Fig.4A-E; Supplementary Fig.5A-D)** compared to the non-injected mice (**Supplementary Fig.5E-H**). However, the level of auto fluorescence was high in these tissues, which range from almost all of the signal (as seen in the heart) to about 40% for 555 nm signals. Thus, in whole section, the analysis of fluorescence level at these wavelengths (555 and 488 nm) may not be relevant in these studies. However, intensity plots show that there were higher levels locally, such as that associated with the airways (bronchioles), liver lobule, kidney tubules and heart tissues (**Fig.4A-D**). The intensity plots of the primary bronchus showed similar distribution as the smaller bronchiole (**Fig. 4F**). Collectively, these data indicate that the CNS vascular barriers restrict the entry of SF-555 and OA-488 but not for peripheral tissues (such as lungs, liver, kidneys and heart). The biodistribution and tissue uptake may be due to ACE2 receptors in these tissues, and non-specific entry. Liver maybe involved in the degradation/elimination of proteins, such as SP. Both of these molecules will be filtered and excreted by the kidneys. SP may be reabsorbed by the kidney tubules, including the proximal tubules [24,25].

**Fig. 4.**
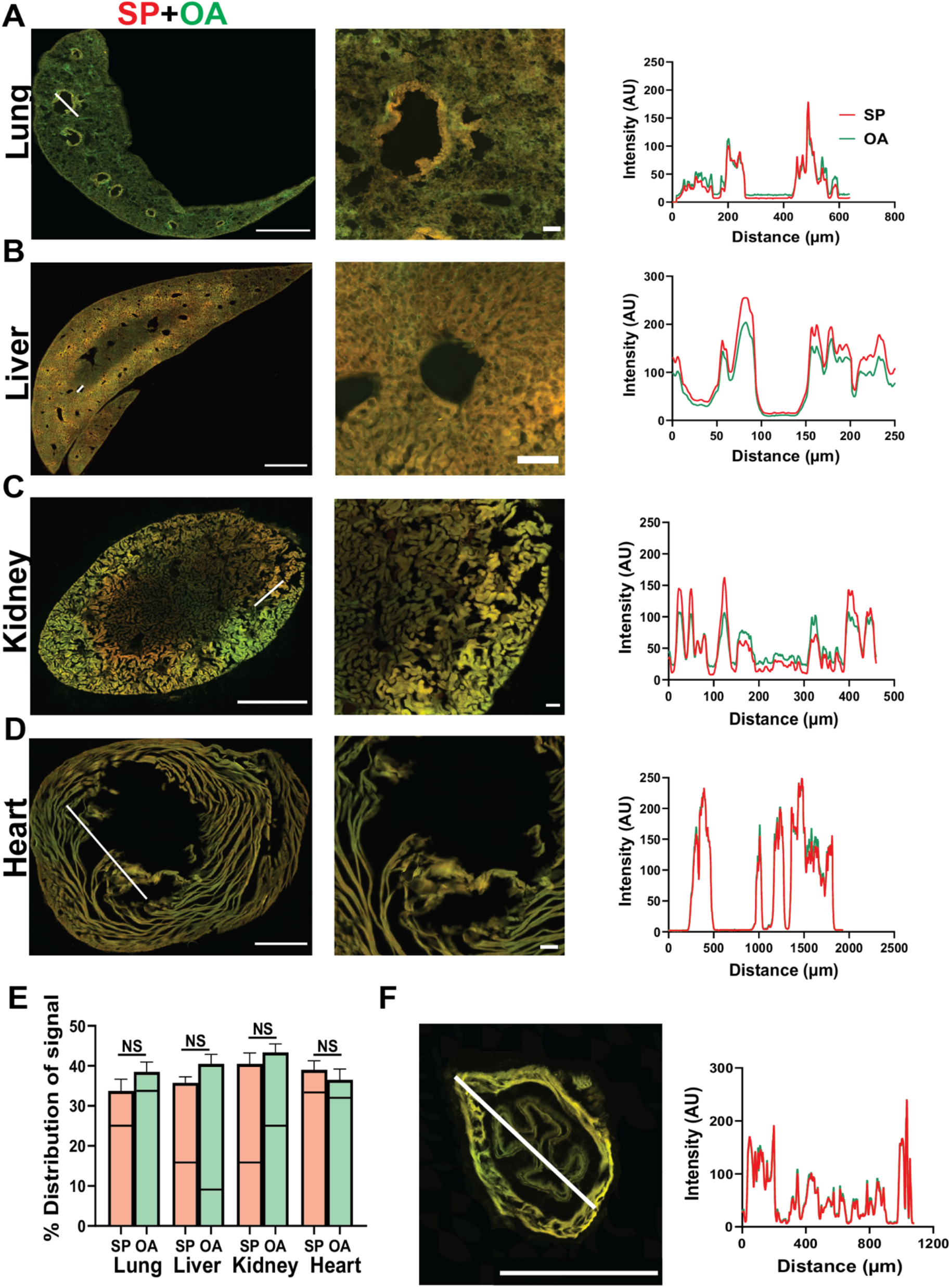
SP-555 peripheral tissues biodistribution. **A-D**) Representative merged images of SP-555 and OA-488 with magnified images and intensity plots at the respective white lines. **E**) Quantification of SP-555 and OA-488 distribution area for the whole section with the auto fluorescence levels (lines) from the non-injected mice. **F**) Merged image of a primary bronchus with intensity plot. Scale bars: A-D and E (1 mm). Values are mean ± SEM, N=3-4 mice.

**Fig. 5.**
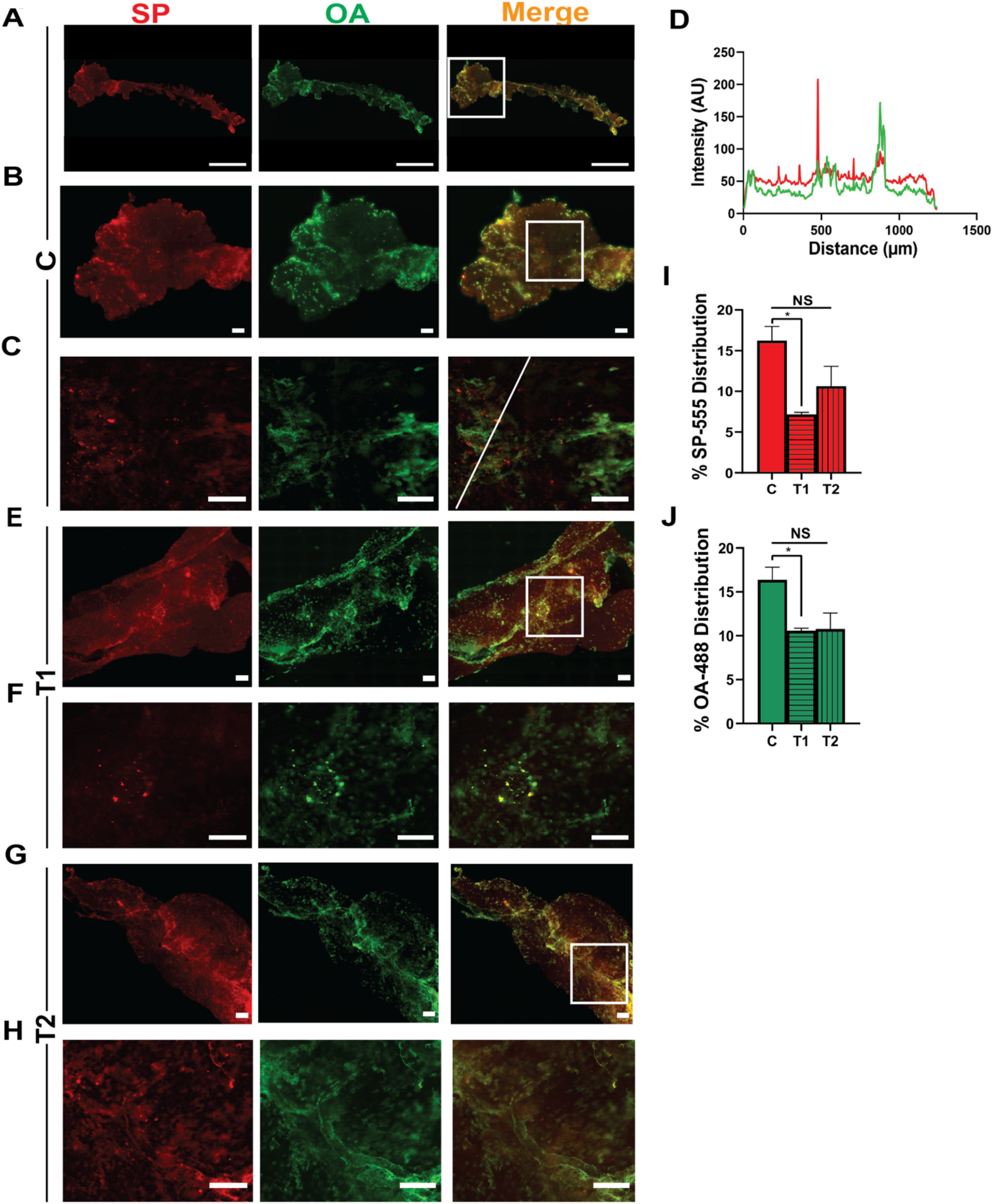
SP-555 is taken up by the isolated CP. **A-C**) Representative images of SP-555, OA-488 and merged images for the isolated CP after incubation for 60 min. Lower panels are the magnified images of the boxed area in the upper panels. **D)** Intensity plot of the line in panel C. It crosses one of the many red dots (SP-555) showing local high intensity. **E-F**) Representative images of SP-555, OA-488 and merged images for the isolated CP after incubation with anti-ACE2 antibody. **G-H**) Anti-SP antibody with the SP-555 and OA-488 for 60 min. Lower panels are the magnified images of the boxed areas. **I-J**) Quantification of the SP-555 and OA-488 signal intensities for the whole areas. Scale bars: A (1 mm); B-H (100 μm). N=3. CP was pre-incubated with T1 and T2 for 15 min and during incubation with the tracers for 60 min.

### SP-555 uptake by the choroid plexus

To confirm the in vivo data on SP uptake by the choroid plexus, the isolated CP was incubated with SP-555 (50 nM) and a reference molecule (OA-488), in the presence and absence of T1 or T2 for 60 min. Images confirmed that SP-555 was associated with the CP (**Fig.5 A-C**). SP-555 location was not always co-localized with OA-488, perhaps due to SP association with epithelial cell and resident leukocytes. Both molecules were present along vessels (**Fig. 5A**). In general, a similar pattern was seen for the in vivo studied, even though the CP was too folded to analyse accurately (**Fig.3E**). Intensity plot across the SP-555 (red) spots revealed high intensity signal (**Fig.5 D**). T1 reduced the association of SP-555 with the choroid plexus, perhaps due to blocking ACE2 receptor binding sites (**Fig.5E-F**). However, T2 was less effective in reducing SP-555 association with the CP (**Fig.5G-J**). This may due to IgG/SP-555 binding to Fc receptors on leukocytes of the choroid plexus [44,45]. Collectively, these data show that SP binds to the choroid plexus, and indicated that T1 altered the SP-555 distribution in the choroid plexus (**Fig. 5 I-J**).

**Supplementary Figure. 1.**
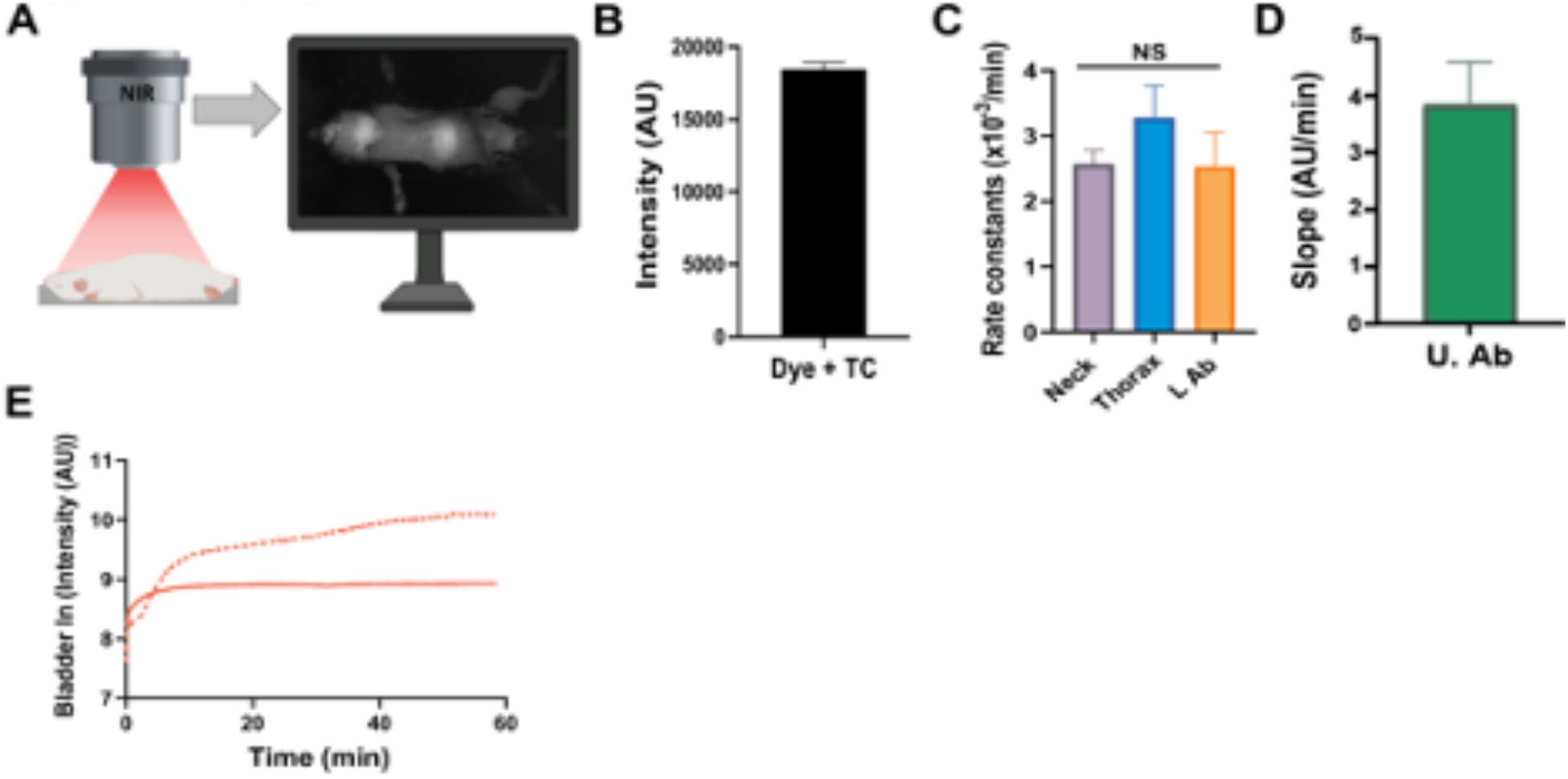
External SP-NIRF in vivo dynamic imaging. **A**) SP-NIRF imaging system for the whole mice. **B**) Confirmation that NIRF signal can be detected from under the isolated mouse rib cage. 10 μL NIRF sample was placed in an Eppendorf tube and covered with the isolated rib cage from a non-injected mouse. **C**) Elimination rate constants for SP-NIRF calculated for ROIs with disappearance part of the profile using the semi- Ln intensity-time profiles. **D**) Slope for the rising phase after SP-NIRF distribution for the upper abdomen. **E**) Variations in urinary bladder SP-NIRF uptake in control mice. Values are mean ± SEM, N=5 young male mice. AU (arbitrary units).

**Supplementary Figure. 2.**
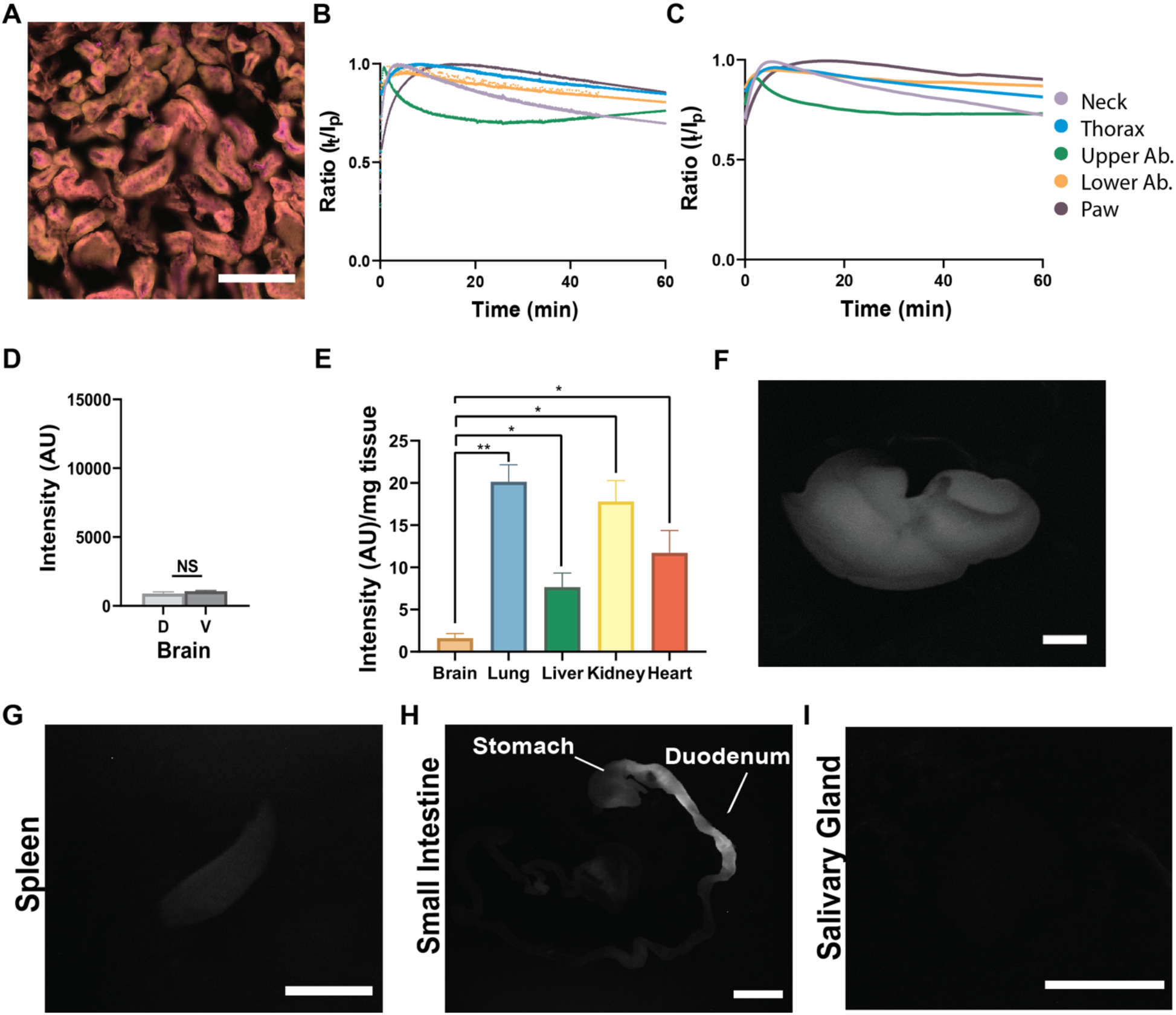
SP-NIRF biodistribution and elimination. **A**) Confirmation of anti-ACE2 antibody-ACE2 interaction in mice kidneys. **B-C**) Standardization of data by dividing intensities at each time point by the peak intensity (IT/IP ratio) for T1 (**B**) and T2 (**C**). **D**) SP-NIRF intensity for the dorsal (D) and ventral (V) brain surfaces. **E**) SP-NIRF intensities/mg wet weight for organs at 60 min. **F-I**) SP-NIRF images for the liver (**F**), spleen (**G**), intestine (**H**) and salivary gland (**I**). Values are mean ± SEM, N=5 young male mice. AU (arbitrary units). Scale bar A (0.3 mm); F-I (1 mm).

**Supplementary Figure. 3.**
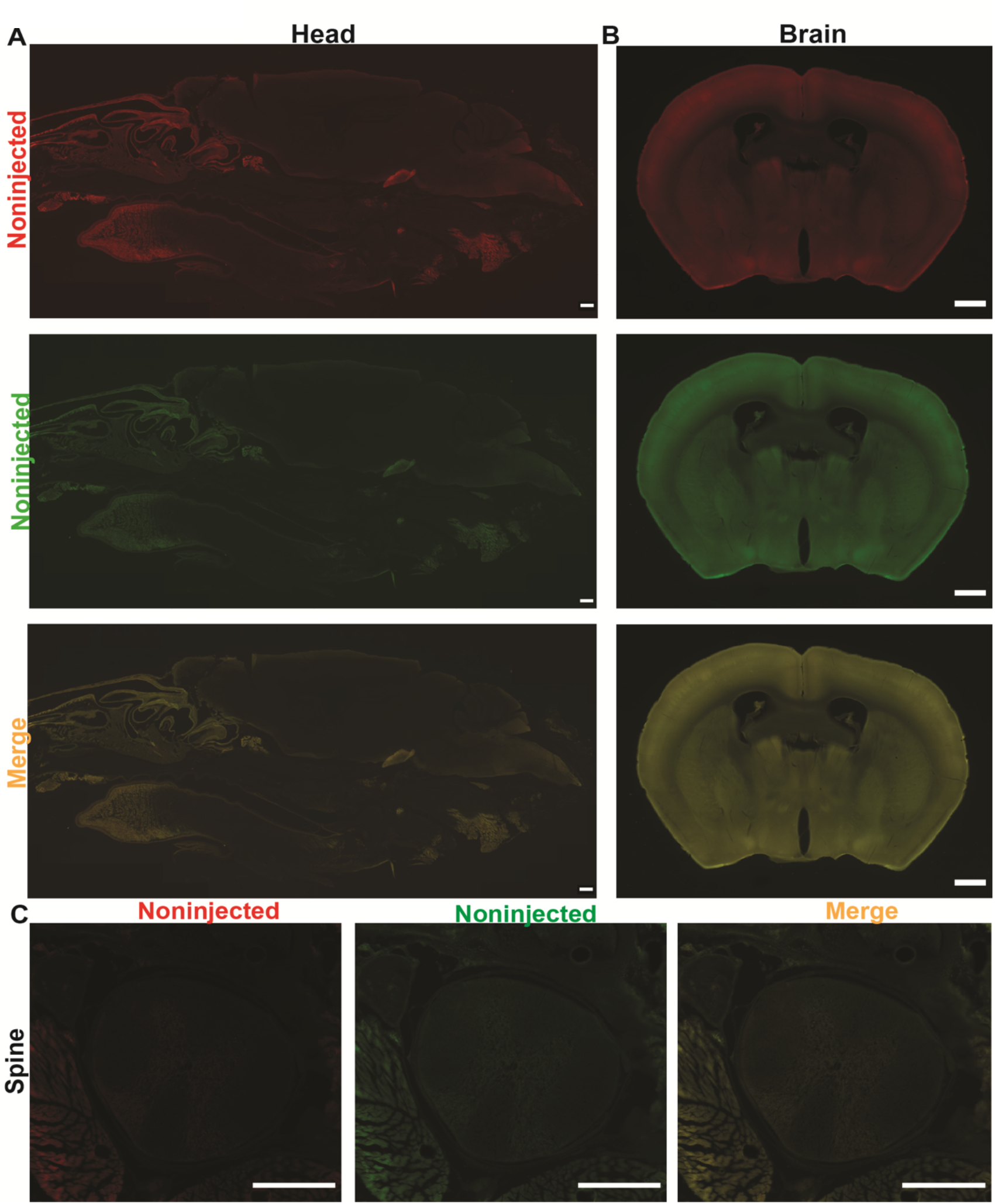
CNS autofluorescence. **A-B**) Representative sagittal section of the head (**A**) and coronal brain section (**B**) for a young mouse showing autofluorescence at wavelengths 555 and 488 nm and merged images. **C**) Representative image of the spine with spinal cord. N=3. Scale bar: A-B (1 mm); C (0.5 mm). Images not enhanced.

**Supplementary Figure. 4.**
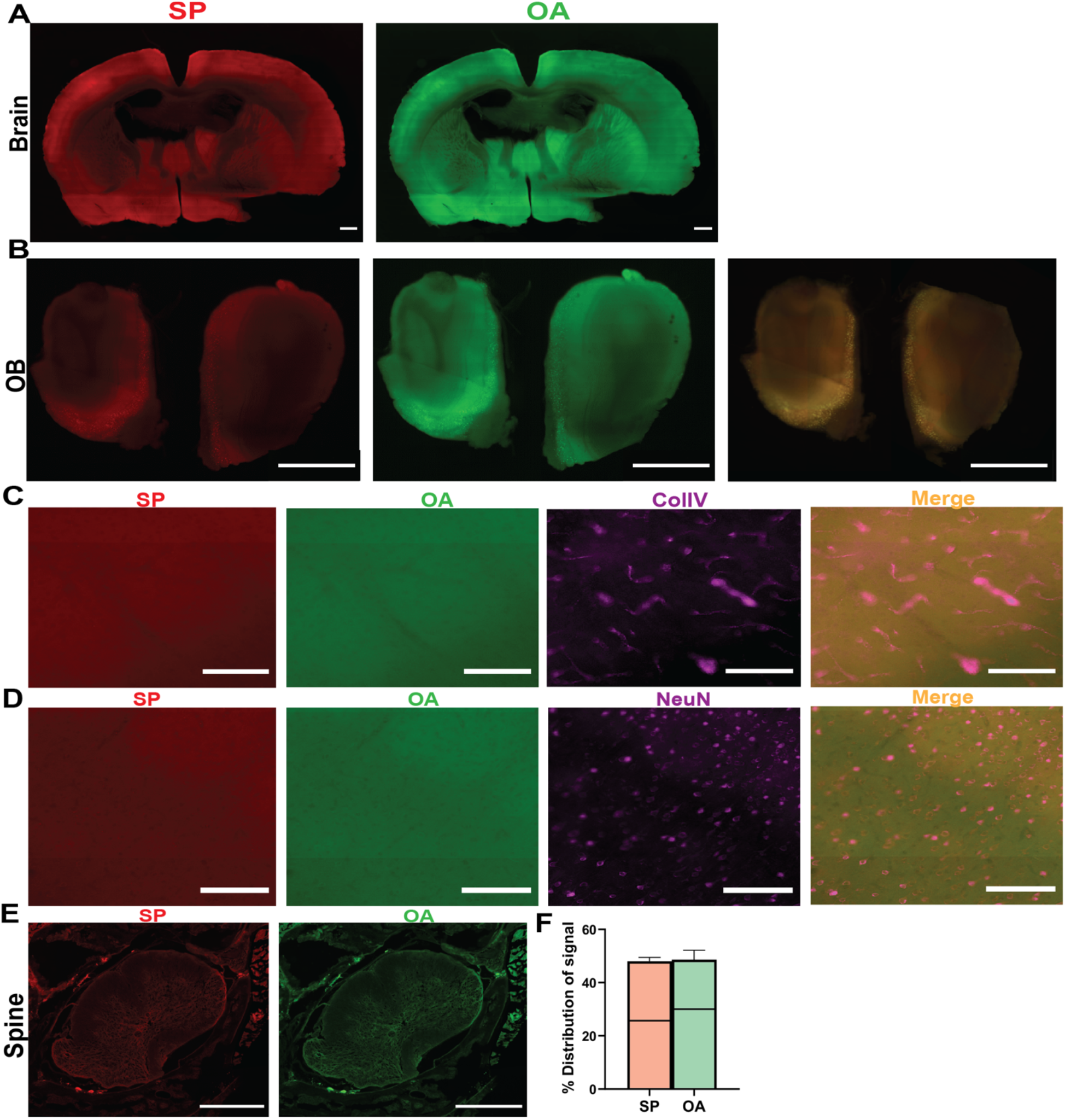
SP-555 is not present in brain parenchyma. **A-B**) Representative coronal sections of brain (**A**) and olfactory bulb (**B**) showing no significant levels of SP-555 or OA-488 compared to autofluorescence. **C-D**) SP-555 in not present in collagen IV-positive vessels (**C**) or NeuN-positive neurons (**D**). **E**) Spinal cord shows some present of the tracers. **F**) Quantification of SP-555 and OA-488 distribution area for the spinal cord only. Lines are the levels of auto fluorescence of the spinal cord from non-injected mice. N=4. Scale bar: A-B (1 mm); C-D (100 μm); E (0.5 mm). Images enhanced for presentation of the details.

**Supplementary Figure. 5.**
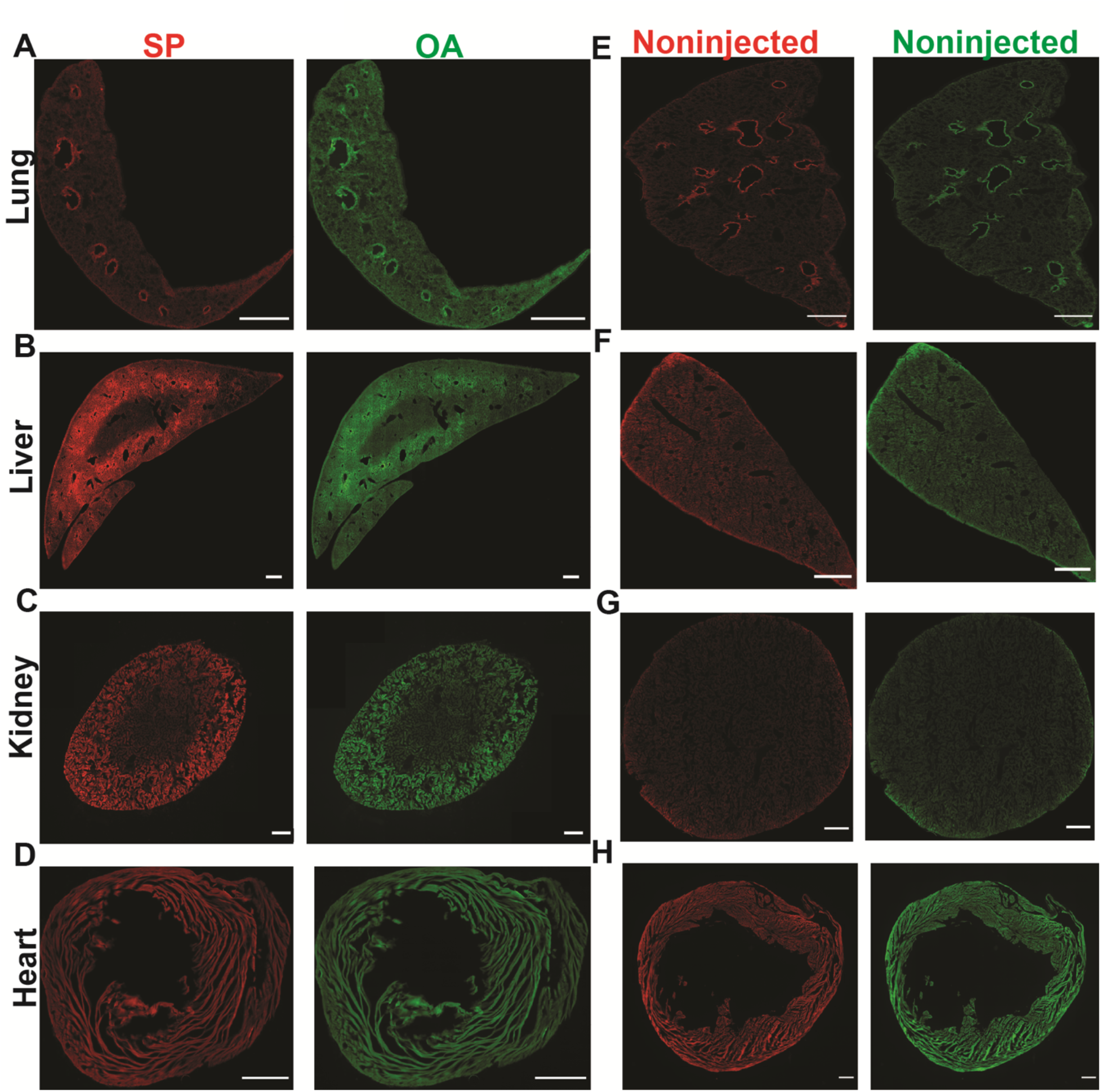
SP-55 and OA-488 are present in peripheral organs. **A-D**) Representative sections (SP and OA) of lung (**A**), liver (**B**), kidney (**C**) and heart (**D**) showing higher levels of SP-555 or OA-488, except for the heart, compared to autofluorescence. **E-H**) Representative images of the lung (**E**), liver (**F**), kidney (**G**) and heart (**H**) showing autofluorescence at wavelengths 555 and 488 nm in non-injected mice. N=3-5. Scale bar: A-H (1 mm).

## Discussion

Our data show that SP-NIRF, administered intravenously in young adult mice (equivalent to young mature humans), was differentially distributed to multiple organs and eliminated slowly from these regions, except the upper abdominal region (mainly liver), which showed a slow accumulation after its distribution. For the paw, SP-NIRF was shown to exponentially increase to a plateau. At 60 minutes post-injection, regional levels were higher than that of plasma, which were undetected. The peak intensities and AUC were highest for the neck region (mainly salivary glands) and upper abdomen, while it was similar for the thorax, lower abdomen and paw. Brain showed the lowest levels, while lung had the highest levels followed by kidney, heart and liver.

While anti-ACE2 and anti-SP antibodies were both effective in reducing SP multiorgan tropism, anti-SP antibody was more effective than anti-ACE2. The brain showed the lowest levels of SP-NIRF signal compared to all organs studied and was not affected by anti-ACE2 or anti-SP antibodies. In contrast, SP-NIRF uptake was significantly reduced with both antibodies for the lung, liver and kidney. There was no significant presence of SP-555 within the brain parenchyma, blood vessels or neuron. However, the choroid plexus was labeled with the injected SP-555, which may contribute to the signal detected in the isolated whole brains, but there was no detectible signal in CSF or the ependymal layer at the interface of CSF and brain. Interestingly, both SP-555 and OA-488 was associated with the spinal cord. Thus, the brain vascular barriers (BBB and choroid plexus) restrict the entry of this viral SP into brain parenchyma, at least within 60 minutes. Our data show that there was differential uptake of SP (lungs highest and brain the lowest) and antibodies against SP or against ACE2 were effective in reducing SP multiorgan uptake, but anti-SP antibody appears to be more effective. By extension, therapeutic antibodies and vaccines that target the SP should minimize the biodistribution and uptake of SARS-CoV-2, and thus, the severity of COVID-19.

We used the receptor-binding domain (RBD) of the SP as a surrogate of the virus (SARS-CoV-2) to study its biodistribution and elimination, since RBD plays a major role for viral entry into host cells. Also, there was no significant differences in the interaction and effects of different sizes of SP once they contain the RBD [32]. In addition, the transfer constant for 2 types of SP1 influx into brain was similar [33,34]. The RBD may have a higher affinity for ACE2 than SP of higher molecular weight [46]. We injected 57 pmol of SP, which were about 34 × 10^12^ monomeric particles using Avogadro’s number. Since there are about 24 SP trimers (72 monomeric forms) on a SARS-CoV-2 molecule [47], an estimated 5 × 10^11^ viruses were injected into blood as SP. Intravenous clearance of viruses was studied at 1.6 × 10^9^ to 1.6 × 10^11^ [48]. Thus, we have used an optimum condition to study SP biodistribution, organ uptake, and elimination.

### SARS-CoV-2

SARS-CoV-2 consists of a viral envelop, which is a lipid membrane that encloses the nucleocapsid bound RNA [49]. The lipid plasma membrane has structural proteins, including the spike protein (SP), which forms a heterodimer (S1-S2) that is assembled as a trimer protruding from the viral surface, which gives the crown-like appearance [10,49] . The S1 unit contains a RBD which promotes attachment by avidly binding to the extracellular peptidase domain on ACE2 receptors on the host cell plasma membranes [10,46,47]. TMPRESS 2 (transmembrane protease, serine 2) on the host cells cleaves the SP to facilitate viral entry [14,50]. Other receptors/facilitators on the surface of host cells have been suggested to mediate the entry of SARS-CoV-2, including sialic acid [51] and CD147 [52].

SARS-CoV-2 enters the body mainly as droplets during inhalation. In critical cases of COVID-19, infection is manifested as cardiopulmonary symptoms, and in severe cases advances into dyspnoea, multiorgan failure and sepsis, which can lead to death in susceptible individuals [1,4,6,53,54]. However, clinical manifestations of the infection (COVID-19) have revealed multiple organs are affected [55]. These non-respiratory organs include the heart [21–23], kidneys [24,25], liver [26–29] liver [26–29], intestine [27,28,56] and brain [3,30,31,34,35,37,57–60]. The distribution of SARS-CoV-2 RNA in a small group of autopsy tissue samples from COVID-19- infected patients shows evidence of multiorgan tropism [55,60].

### ACE2/SARS-CoV-2

ACE2 is expressed on the cell membrane in many tissues, including the gastrointestinal tract, kidneys, choroid plexus and bladder [10,13,14]. It is expressed in the heart, vascular endothelial cells, and vascular smooth muscles cells of arteries and venules [61]. ACE2 is highly expressed on type 1 and type 11 epithelial cells and bronchiolar epithelium [11,15]. In addition, it is also expressed on the oral mucosa [41]. In the brain, ACE2 mRNA is present in the cortex, striatum, hippocampus and brain stem [16–18]. It is mainly expressed in the cytoplasm of neurons, and in brain regions associated with regulation of the cardiovascular system (CVS), blood pressure (BP) and the autonomic nervous system [19]. It was reported that the brain endothelium expresses ACE2 as the protein [32] and as the RNA-seq [11,62]. The brain vasculature/vascular endothelium shows a high expression of the SARS-CoV-2 protease cathepsin B (CTSB) [42]. ACE2 is highly expressed on pericytes [12]. While there are conflicting reports on ACE2 distribution and its significance to an individual organ, the wide expression of ACE2 receptor suggests that many organs will be affected by SARS-CoV-2 to some degree. This may be due to direct interaction with ACE2 receptors of a specific organ or indirectly as a consequence of cardio-respiratory failure. The precise location (extracellular/intracellular) of ACE2 in the endothelium is needed for target organs.

### SP biodistribution from blood to brain

The mammalian CNS is unique in that it is enclosed within its vascular barriers, and has cerebrospinal fluid (CSF) continuously circulating around it [63–66]. The vascular barriers at the blood-brain interface (the blood brain barrier, BBB) and blood-CSF interface (blood CSF barrier; the choroid plexuses) restrict the diffusion of polar molecules and large molecules into and out of the brain, while essential metabolites are transported into brain and/or into CSF [63–66]. The physical sites of the BBB and blood CSF barrier are the tight junctions between endothelial cells and epithelial cells, respectively [67,68], which restricts paracellular diffusion. The endothelium of the cerebrovasculature is at the interface between the blood and brain, but there are several cell types on the abluminal side of the endothelium. These include pericytes, astrocyte end-feet and microglia [63,64]. The SP will need to interact with the luminal surface of the endothelium to enter brain from blood.

CSF is mainly produced within the cerebral ventricles by the choroid plexuses, and circulate from the lateral ventricles to the subarachnoid space, around the brain, spinal cord and within the spinal canal, before ultimately draining at multiple outflow sites into blood [65,69–73]. A major CSF drainage pathway is via the olfactory bulb, across the cribriform plate and towards the cervical lymphatic nodes, and the spinal cord [39,65,73]. The endothelium of the choroid plexus is fenestrated, and allows entry of proteins into the stroma, which is restricted from entering CSF due to the tight junctions between the epithelial cells at the apical surface [63,65,66].

Following intravenous SP-NIRF injection, it will be distributed to all organs via blood, and uptake determined by the degree of restriction offered by the endothelium of the vasculature [91,92]. Continuous endothelium of the CNS vascular barriers offers the greatest restriction to protein flux, except at the CP where the endothelium is fenestrated and allow a greater flux into the stroma [63–66]. Thus, both SP-555 and OA-488 will cross the fenestrated endothelium of the CP, but not the epithelium layer, which has tight junctions at its apical side between these cells [65].

Our data confirm multiorgan tropism seen in clinical manifestations of COVID-19, with the exception of brain. While the CP was labeled with SP-555, there was no detection of SP-555 in CSF, since the epithelium, likely, restricts its entry. However, there was SP distribution to the spinal cord, which may reflect the greater permeability of the blood-spinal barrier to large (albumin) and small (inulin) molecules compared to brain [43]. Thus, the brain vascular barriers at the BBB and CP seem to restrict the entry of SP into brain and CSF from blood. There is a report that SP interacts with the brain endothelial cells using in vitro models of the BBB [32]. In addition, there is a report that SP enters the murine brain from blood [33,34]. Like this report, we found that the SP-NIRF brain levels were not dependent on ACE2. The time point for these reports and our study was similar. In our study, the low levels of SP-NIRF in brain was likely associated with the CP. Thus, our data support the finding that CP is a potential target of SARS-COV-2. SARS-CoV- 2 was shown to be associated with the CP and can damage the epithelium [74] and SARS-CoV-2 transcripts were associated with the CP [58]. In addition, there are reports both for the presence of SARS-CoV-2 in the CSF [5,57] and other reports that it is not in CSF [75,76]. We found that SP- 555 also binds to the apical surface of the CP using the isolated CP from mice, which may also limit SARS-CoV-2 in CSF. This may be due to ACE2, but the highly convoluted apical surface of the CP and its folding make it difficult to analyze tracer distribution. However, there are spots of high intensity SP-555 not associated with OA-488. Further work is needed to identify these cells that selectively take up SP-555.

The exact mechanism of SARS-CoV-2 neuroinvasion is unclear. Viruses could enter the brain by retrograde transport via sensory nerve endings within the nasal and buccal cavities, such as the cranial olfactory and trigeminal, and the autonomic nervous system [77–80]. There is also the possibility of viral neuroinvasion via the gastrointestinal tract [57,81].

It is possible that SARS-CoV-2 enters brain as a consequence of pneumonia induced-hypoxia. However, there are reports of encephalitis, which was not due to COVID-19-induced hypoxia [82] and brain cortical hyper-intensity, as seen in MRI images, which may be due to viral infection [3,83]. The virus was detected in frontal lobe brain tissue, which may suggest access via the nasal route [84]. There are also reports of the presence of SARS-CoV-2 in autopsy brain samples and especially in the olfactory mucosal region of severely infected patients [78–80,85].

However, this is not always the case, as SARS-CoV-2 RNA was not detected in the nasopharyngeal swab of a patient but was detected in CSF [5]. The virus RNA was detected in the CSF of a COVID-19 patient with meningitis [3]. Viral particles are associated with the cerebrovasculature and the endothelium of other organs [86–90]. However, it’s unclear if these clinical neurological presentations seen in severe COVID-19 patients are due to the virus entering brain or as a consequence of cardio-respiratory, multiorgan failure and systemic inflammation. It is possible that the viral invasion of the lungs and the subsequent inflammation will dominate and drive the outcomes of the infection. Thus, further studies are needed to support significant neuroinvasion as the explanation for the neurological symptoms associated with COVID-19.

### SP biodistribution from blood to peripheral organs

We performed *in vivo* dynamic imaging in regions (ROIs) that were known to be affected by COVID-19. While these ROIs are representing multiple tissues, in some of them signals were associated with a predominant organ. These included the neck region where the SP-NIRF signal was predominantly from the salivary gland and the upper abdomen where it was from the liver. For the thoracic ROI, the NIRF signal was most likely from the lungs and heart, while the signals from the lower abdomen are from mainly the intestine and kidneys. Signals from the paw are due to the hairless skin.

There is a continuous endothelium with tight junction between endothelial cell for the lung which should limit protein flux [93–95]. However, unlike the brain, the lungs had the highest SP- NIRF uptake from blood. This may be due to the heterogeneity of the endothelial cells and SP- NIRF binding to the glycocalyx and ACE2 receptor [92,95] For the liver, the sinusoidal endothelium and Kupffer cells clear viruses and proteins [48,96–99]. ACE2 is expressed on many parts of the nephron, but especially on the luminal brush border of the proximal tubule [100–103]. Thus, the kidney likely takes up a significant level of SP. Also, ACE2 is expressed in the heart [21–23,102,104]. Our findings that SP is distributed to the salivary glands, may support suggestions that these glands are a potential reservoir of SAR-CoV-2, which may lead to parotitis- like syndrome [105,106]. Interestingly, within the gastro-intestinal tract, blood-borne SP-NIRF was mostly located at the duodenum [27,28,56,107,108]. The intestine expresses ACE2 and this has been suggested to contribute to COVID-19-related gastrointestinal tract effects [27,28,56,108]. However, it unclears on how much of the ACE2 is intracellular and extracellular in all tissues. In addition, it’s unclear on whether expression is translated to protein levels, and more specifically whether ACE2 protein is extracellular on the endothelium cell membranes.

### Role of anti-ACE2 and anti-SP antibodies in SP biodistribution

While the antibodies were injected 15 mins prior to the tracer injection, they will remain in the circulation for a longer time as the half-life is very long, about 18 to 20 minutes [109]. Receptor-mediated endocytosis of IgG by FcRn-mediated recycling of IgG molecules may contribute to this process. In mice, many organs are involved in IgG clearance, including the liver, kidney, muscle, skin and spleen [109,110]. The rationale for the current study is that for the anti-ACE2 antibody, it will interact with ACE2 receptor and reduce SP binding to the host cells [111]. In contrast, anti-SP antibody will interact with the SP and reduce the free SP levels, which in turn will reduce SP binding to tissues. Our data show that the anti-SP antibody was more effective in reducing SP biodistribution and uptake into organs [112]. The anti-SP antibody was more effective in reducing SP-NIRF uptake than that of anti-ACE2 antibody in lungs, liver and kidneys. In contrast, there was no effect of these antibodies on brain SP-NIRF uptake. It’s possible that IgG-SP could bind to Fc receptors on cells, especially leukocytes. The CP has resident immune cells, including, macrophages and CD4 cells [44,45], which could have a greater effect on SP-NIRF uptake by the isolated CP, which may not be seen *in vivo* as immune cell will eliminate these.

### Limitations of the present study

We used the RBD of the SP as a surrogate of the virus (SARS-CoV-2) to study its biodistribution and elimination, since RBD plays a major role for viral entry into host cells. However, the distribution pattern might be different for the actual virus, and the molecular weight of SP would influence filtration at the kidneys and convective flow in peripheral organs, such as muscles. Thus, a lower molecular weight RBD (< albumin) likely represent a maximum biodistribution. The biodistribution of SP was not systematically evaluated in all organs/cells, such blood cells, pancreas and others. Therefore, it is possible that an organ not studied may also play a role in SP uptake. However, these other organs may not be as significant, since there was no significant SP- NIRF level in the rest of the mouse. Thus, we focused on organs with significant signal. The interaction of the whole virus with host cells may be different. Nevertheless, our studies and other that use the SP would provide critical data to better and safely model the virus behavior in a host before using the virus in studies. In the current study the efficacy of anti-SP and anti-ACE2 antibodies was assessed and thus, intravenous injection of SP was used. However, the main route of viral entry into the body in inhalation of droplets. In terrestrial animals, injection of fluid into the nose is not physiological. The SP would need to specially formulate into an aerosol for inhalation studies, which is beyond the scope of this study. In addition, high levels would be needed to enter blood via inhalation to meet the objectives of the current study. In patients, blood levels of SARS-CoV-2 is low [55]. Plasma profile of SP was not possible as serial sampling of large volume of plasma would be needed. The total blood volume is ∼ 1.5 ml (in a 20 g mouse and assuming blood volume is 78 ml/kg body weight (vivarium)) and plasma volume ∼0.9 ml. However, the same dose was administered intravenously and all mice were of the same age and sex.

## Conclusions

SP-NIRF is differentially distributed to many organs, which may explain SARS-CoV-2 multiorgan tropism, and this may contribute to organ failure, in addition to respiratory failure, in young adults. The lungs had the highest SP levels and the brain the lowest. There was significant SP signal in the salivary glands, intestine, spinal cord and choroid plexus, but not in CSF. Thus, the brains vascular barriers were effective in restricting the flux of proteins from blood into the brain parenchyma. The differential SP organ uptake is likely determined by, mainly, ACE2 levels. Anti-SP antibody was more effective than anti- ACE2 antibody in suppressing SP biodistribution and organ uptake. Thus, therapies, include passive immunity using anti-SARS-CoV-2 antibodies, and convalescent plasma, which contains anti-SARS-CoV-2 antibodies, will be effective in reducing SARS-CoV-2 biodistribution, and thus COVID-19 severity. It also offers confirmation that vaccines against the SP, which generates anti-SP antibodies, are like to neutralize the virus tissue distribution and minimize severity of this infection. Further work is needed in older mice and with systemic inflammation.

## Material and Methods

### Materials

SARS-CoV-2 Spike proteins (recombinant SARS-CoV-2 Spike Protein (S-RBD; cat# RP-87678, HEK293 cell expressed and binds ACE2) and ovalbumin Alexa Fluor-488 conjugate (Cat# 034781; molecular weight 45 kDa) were obtained from Life Technologies Corporation, Carlsbad CA, USA. The spike protein (molecular weight 36 kDa)) was labeled with NIRF using a kit (IRDye800CW Microscale kit, Li-COR Biosciences, Nebraska, USA) by following the manufacturer instructions. This conjugation forms a stable amide bond, and has been used in clinical trials (Li-COR Biosciences). SP was labeled separately with Alexa Fluor 555 using a kit (Microscale protein labeling kit; ThermoFisher Scientific; Waltham, MA, USA) and by following the manufacturer instructions. In addition, both labels were purified using 3 kDa molecular weight cut-off ultrafiltration filter (Amicon Ultra Centrifugal Filter, Millipore). There was no detectable dye in the filtrate. Anti-SARS-CoV-2 spike protein antibody (SARS-CoV-2 Spike Protein Monoclonal Antibody (cat# MA5-36087; immunogen SARS-CoV-2 spike protein that interact with HeLa cell expressed SARS-CoV-2) and anti ACE2 antibody (ACE2 Recombinant Rabbit Monoclonal Antibody (Cat# MA532307; interact with mouse ACE2) were obtained from Life Technologies Corporation.

### Mice

C57BL/6J (2-3 months old; Jackson Laboratory; Bar Harbor, ME, USA), male mice were used. All animal studies were performed in accordance with the National Institute of Health guidelines and using protocols approved by the University Committee on Animal Resources. Mice were housed in the vivarium of the University of Rochester, School of Medicine and Dentistry on a 12:12 light/dark schedule (6AM: 6 PM) with food and water ad libitum.

### Intravenous Injections

Mice were anesthetized with isoflurane since it has fewer systemic hemodynamic effects [38] and placed on a temperature-controlled stage to maintain body temperature. Anesthesia was maintained with 1-2% isoflurane in oxygen. Once anesthetized, a midline incision from the neck to the pelvic area was made, and the skin retraced to enhance the NIRF signal intensity. Local anesthetic (Topical Lidocaine 4% gel, ESBA Laboratory Inc., FL, USA) was applied to the exposed regions, which was moistened with saline. This also exposed the right external jugular vein for a minimal invasive intravenous injection using a 30 G needle connected to a 500 µL insulin syringe via a tubing. The tubing contained the fluorescent tracer (10 μL), which was the spike protein conjugated to IRDye800CW (SP-NIRF; 5.7 pmoles/μL), or spike protein conjugated to Alexa-555 (SP-555; 5.7 pmol/μL) and a protein reference molecule of similar molecular weight (ovalbumin-488 (OA-488, 0.1%), dissolved in PBS (pH 7.4). The injection contained 10 μL PBS, 1 μL air gap, 10 μL tracer, 1 μL air gap, and 50 μL PBS to wash in the tracers completely. The injection was completed in 1 minute. We used NIRF to minimize the natural background fluorescence of biomolecules, increase the signal to noise signals, and to provide a better contrast between the target and background (e.g., [39]). We also used SP-555 to visualize and map its biodistribution in tissue sections, and to compared this to that of a reference protein molecule of similar molecular weight (OA-488).

The groups of mice were as follows: young male mice (2-3 months old), T1 (anti-ACE2 antibody; young male mice (2-3 months old)) and T2 (anti-spike protein antibody; young male mice (2-3 months old)). T1 (10 μg) or T2 (10 μg) was injected instead of the 10 μL PBS 15 min prior to the tracer (the same dose). The experimenter was blinded to the experiment design, and unaware of the tracers, T1 and T2 used in these studies.

### In vivo dynamic imaging

NIRF intensities were measured using a custom-made NIR system (**Fig. S1A**). Mice were placed on a heated surface (Indus Instruments, Webster, TX, USA), and the hair was removed with a depilatory cream before surgery. The imaging system was composed of a lens (Zoom 7000, Navitar, Rochester, NY, USA), a NIRF filter set (Semrock ICG-B, IDEX Health & Science LLC Rochester NY) and camera (Prosilica GT1380, Allied Vision Technologies, Exton, PA, USA). NIRF was excited with a tungsten halogen bulb (IT 9596ER, Illumination Technologies, Inc., Syracuse, NY, USA) through a ring illuminator (Schott, Elmsford, NY, USA). Imaging settings and recordings were accomplished through a custom-built LabVIEW program (National Instruments, Austin, TX, USA). Real time NIR imaging was performed before injections (background) and every 2 seconds for 60 minutes after SP-NIRF injection into the right jugular vein to quantify its biodistribution [39,40]. Using ImageJ software (National Institutes of Health, Bethesda, MD, USA), regions of interest (ROIs) were identified, and the ROI fluorescence intensity recorded at the same settings, as reported [39]. The person analyzing the data were blinded to the experimental design.

### Ex vivo NRIF imaging of whole tissues

At the end of in vivo imaging, animals were sacrificed and tissue samples harvested, including brain, spinal column, liver, lungs, kidneys, heart, spleen and intestine. Tissues were washed equally and imaged individually on the ventral and dorsal surfaces using the same parameters as for in vivo imaging. The average tracer intensities were used as there was no significant difference in the dorsal and ventral values.

### Tissue Preparation for fluorescence imaging

After the duration of the experiment, mice were transcardially perfused using cold phosphate buffered saline (PBS) and paraformaldehyde (PFA) (4% in PBS, pH: 7.3). Tissues samples were removed, stored overnight in PFA at 4 °C and re-stored in cold PBS. Brain was cut into 100 µm coronal sections using a vibratome (Leica VT1000E). Spinal columns (the cervical region removed), liver, kidney, lungs and heart were embedded in Optimal Cutting Temperature (OCT) compound and cut into 30 μm sections using a cryostat. In some experiments, the skin was removed from the head, then decalcified in 13% EDTA for 5 days, and cut using a cryostat. Sections were mounted on Superfrost Plus glass slides using ProLong Gold Antifade Mountant medium (ThermoFisher Scientific; Waltham, MA, USA) for fluorescence imaging (VS120 Virtual Slide Microscope, Olympus). Exposure and gain were fixed for all experimental groups based on pilot experiments. All fluorescence (SP-555 and OA-488) quantification was performed without enhancement of signals. This was finally decoded when the figures were prepared for publication.

### Ex vivo Fluorescence Imaging

Ex vivo epifluorescence microscopy was used to evaluate the degree of tracer distribution in the tissue sections. Exposure and gain were fixed for all experimental groups based on the pilot experiments. To quantify the extent of tracer intensity distribution in whole tissue sections, the whole-slice/section images were analyzed using ImageJ (National Institutes of Health, imagej.nih.gov/ij/), and Qupath (https://qupath.github.io/). In Qupath, ROI was created for each analyzed sample. A training image was created from the (ROIs) to train a three-way random trees pixel classifier to account for variation, and distinguish between positive fluorescence and negative background. Separate pixel classifiers were created for 555 and 488 nm channels. For each slice, fluorescence emission channels were split into individual channels of 8-bit TIFF grayscale images and a whole-section ROI was defined. The pixel area of positive fluorescence (threshold pixel intensity for both green (488 nm) and red (555 nm) channel was calculated for each section and the average expressed as a percentage of the whole section area. The Olympus VS120 Virtual Slide Microscopy was used to identify co-localization of immunolabelled (cellular or vascular components) with the injected tracer.

### Kinetics analysis

For each experiment, image sequences were imported into ImageJ. The ROIs, identified from pilot experiments were neck (including salivary glands, trachea, pharynx), thorax (lungs, heart), upper abdomen (including liver, spleen), lower abdomen (intestine, kidneys but not bladder) and paw (a type of skin region with no clear relationship with COVID-19) (**Fig. 1A**). Mean pixel intensity within each ROI was measured for each time point using the images without enhancement. The background signal was the same ROI without the injection and this was subtracted from the intensity at each time point. To correct for experimental variation, the intensity at each time point was divided by peak intensity to standardize the data. Peak intensity (Imax) was identified as the average of 10 seconds at the peak identified using GraphPad Prism software (Version 9.1.0 (216)). The area under the curve (AUC) was determined by running AUC analysis option in GraphPad Prism. The elimination rate constant was determined from the slope of the falling phase of the intensity profile using a semi-natural logarithmic (Ln) plot and the linear regression function in the GraphPad Prism. For the liver, since there was a linear component of the intensity-time profile after the distribution phase, the slopes were determined from the plots using GraphPad Prism.

### CSF and blood samples

In a separate group of experiments, after tracer injection (young mice) CSF and blood samples were collected. CSF was collected from the cannulated cisterna magna (to minimize contamination from blood) and samples from two mice pooled to get a volume to analyze (15 μL). Blood samples was collected by cardiac puncture (100 μL). A fixed volume (15 μL) of plasma and CSF in an Eppendorf tube was used to determine NIRF intensities and for comparison. Background was the same volume of saline an Eppendorf tube. The total volume of plasma in a mouse is < 1 ml.

### Choroid plexus uptake

In a separate group of experiments, young mice were perfused with cold PBS and the choroid plexuses (CP) isolated from the lateral ventricles. These were incubated in 100 μL of oxygenated (100% O2) HBSS, and in a humidified oxygenated chamber at 25 ^0^C for 60 minutes. There were three groups, control, T1 and T2. The traces were 50 nM SP-555 and the reference molecule OA-488 (0.01%). T1 or T2 (100 nM) were added 15 minutes before the tracers. At the end of the experiment, the incubation media were removed, CP washed 3 times with 500 μL cold PBS, fixed in 100 μL 4% PFA, washed 3 times with 500 μL cold PBS, mounted on glass slides and imaged.

### Statistical analysis

Data were analyzed by analysis of variance (ANOVA) followed by post hoc Tukey test using GraphPad Prism. Differences were considered to be significant at p < 0.05. For statistical representation, P* <0.05, P** <0.01, and P*** < 0.001 and P**** p <0.0001 are the levels of statistical significance. NS is not significant. All values were expressed as mean ± SEM. Student’s t-test with Welch’s correction was used for sample comparison. GraphPad Prism software (version 9.1.0 (216)) was used for all analysis.

## Acknowledgements

This work was supported by NIH grants to RD (AG057574; R21AG050212) and to the Center for Musculoskeletal Research Histocore (CMSR) core facility (NIH P30 NIAMS AR069655).

